# SAN: mitigating spatial covariance heterogeneity in cortical thickness data collected from multiple scanners or sites

**DOI:** 10.1101/2023.12.04.569619

**Authors:** Rongqian Zhang, Linxi Chen, Lindsay D. Oliver, Aristotle N. Voineskos, Jun Young Park

## Abstract

In neuroimaging studies, combining data collected from multiple study sites or scanners is becoming common to increase the reproducibility of scientific discoveries. At the same time, unwanted variations arise by using different scanners (inter-scanner biases), which need to be corrected before downstream analyses to facilitate replicable research and prevent spurious findings. While statistical harmonization methods such as ComBat have become popular in mitigating inter-scanner biases in neuroimaging, recent methodological advances have shown that harmonizing heterogeneous covariances results in higher data quality. In vertex-level cortical thickness data, heterogeneity in spatial autocorrelation is a critical factor that affects covariance heterogeneity. Our work proposes a new statistical harmonization method called SAN (Spatial Autocorrelation Normalization) that preserves homogeneous covariance vertex-level cortical thickness data across different scanners. We use an explicit Gaussian process to characterize scanner-invariant and scanner-specific variations to reconstruct spatially homogeneous data across scanners. SAN is computationally feasible, and it easily allows the integration of existing harmonization methods. We demonstrate the utility of the proposed method using cortical thickness data from the Social Processes Initiative in the Neurobiology of the Schizophrenia(s) (SPINS) study. SAN is publicly available as an R package.

## 1 Introduction

Emerging techniques facilitate the quantification of human cerebral cortex properties through magnetic resonance imaging (MRI), such as cortical thickness, surface area, and gyrification (Goto et al., 2022). These advancements have profound implications in the study of brain structures and functions. Cortical thickness is defined as the spatial span between the gray matter and white matter surfaces of the cerebral cortex, for which subtle variations in brain structure may enhance the understanding of neurological disorders, developmental changes, and potential cognitive processes.

Notably, alterations in cortical thickness have been implicated in normal aging (Thambisetty et al., 2010; Sowell et al., 2003), as well as conditions such as Alzheimer’s disease (Cho et al., 2013; Lerch et al., 2005), schizophrenia (Van Haren et al., 2011; Narr et al., 2005), and multiple sclerosis (Sailer et al., 2003; Tsagkas et al., 2020).

There are two main approaches for analyzing cortical thickness data: the region-level analysis and the whole-brain analysis (vertex-level). The region-level analysis first uses a brain parcellation atlas (e.g., Desikan-Killiany atlas) and obtains averaged cortical thickness data for each region of interest (ROI) (Hanford et al., 2016; Li et al., 2022). It offers the simplicity of achieving dimension reduction and alleviating multiple comparison problems better than whole-brain univariate analysis. However, it is limited to predefined regions, so it is unable to localize ‘signal clusters’ (patterns associated with certain conditions) that might emerge in smaller areas within ROIs or that span across multiple ROIs. Another approach is whole-brain analysis to analyze cortical thickness from all vertices throughout the brain (Goto et al., 2022; Bernal-Rusiel, Greve, et al., 2013; Ge et al., 2015), which offers better localization of signals at the expense of an increased burden in multiple comparisons (Lindquist & Mejia, 2015). On a positive note, recent studies on leveraging spatial dependencies inherent in vertex-level cortical thickness data have led to high power and effectively controlled the false positives (Bernal-Rusiel, Reuter, et al., 2013; Park et al., 2021; Weinstein et al., 2022). Moreover, spatial covariance modelling in neuroimaging has shown evidence for promising performance in other neuroimaging modalities when integrated with cluster enhancement (Park & Fiecas, 2022; Pan et al., 2024).

Large-scale neuroimaging studies often use multi-site, multi-scanner protocols to recruit study participants quickly and in large numbers. However, a major challenge of combining neuroimaging studies across sites/scanners is *inter-scanner biases* that are introduced due to several technical variabilities in these studies, including disparities in scanner manufacturers, variations in scanner parameters, and heterogeneities in acquisition protocols (Han et al., 2006; Schnack et al., 2010; Jovicich et al., 2006). As with other imaging modalities, these inter-scanner biases have been shown to be present in vertex-level cortical thickness data (Schnack et al., 2010), which motivates a need for addressing inter-scanner biases and providing high-quality cortical thickness data for downstream whole-brain analysis.

Several statistical harmonization methods have been developed to identify and parameterize the source of inter-scanner biases and mitigate them by reconstructing new homogenized data for downstream analysis. One prominent approach, ComBat (Johnson et al., 2007), first proposed in genomics, has been adapted for the removal of inter-scanner biases across various neuroimaging data modalities, including DTI mean diffusivity and fractional anisotropy (Fortin et al., 2017), region-level cortical thickness (Fortin et al., 2018), and functional connectivity (Yu et al., 2018). ComBat characterizes scanner effects into an additive (mean) and a multiplicative (variance) scanner effect for each imaging feature. Moreover, ComBat has been extended to harmonize imaging data collected in a longitudinal manner (Beer et al., 2020) and to harmonize MRI scans at the voxel level by incorporating the superpixel technique (C.-L. Chen et al., 2022).

Recent harmonization methods (CovBat and RELIEF) have shifted to expand the scope of statistical harmonization to address heterogeneous covariances, going beyond the mean-variance specifications in ComBat (A. A. Chen et al., 2022; Zhang et al., 2023). These methods have shown that successful harmonization of covariances improves signal-to-noise ratios, thus preserving biological variations better. However, their existing applications have primarily focused on region-level data, and there is limited empirical evidence to assess their efficacy for vertex-level data. Also, while these methods are generally applicable to any neuroimaging data types (which supports their versatility), at the same time they are expected to perform suboptimally when there exist specific covariance patterns that cannot be addressed solely by data itself. For example, the low-rank decomposition used by both CovBat and RELIEF lose rich local spatial information, which would be suboptimal for vertex-level cortical thickness data with a significant degree of spatial autocorrelation. Furthermore, existing harmonization methods do not preserve the spatial smoothness of the harmonized data, which raises a critical issue when they are used in downstream analysis with spatial covariance modelling.

To address these challenges, we propose a new harmonization method called Spatial Autocorrelation Normalization (SAN) to identify and parameterize the sources of inter-scanner biases in vertex-level cortical thickness data and reconstruct homogenized and spatially smooth data. A central challenge of SAN lies in modelling scanner-specific spatial covariances, differentiating them into heterogeneous non-spatial variations and spatial variations, all the while preserving the underlying homogeneous autocorrelation structures across scanners. In SAN, we use the spatial Gaussian process to leverage pairwise distance in the characterization of covariance heterogeneity, which effectively addresses the local patterns of covariance heterogeneity and preserves spatial smoothness in the data. We apply our method to the Social Processes Initiative in the Neurobiology of the Schizophrenia(s) (SPINS) study, a multi-site, multi-scanner neuroimaging study including participants with schizophrenia spectrum disorders and healthy controls, to validate SAN, then conduct a data-driven simulation to compare SAN to other harmonization methods.

## 2 Methods

### 2.1 Notations and model specifications

#### 2.1.1 Characterization of heterogeneous means and variances

We let *y*_*ijv*_ be an imaging feature measured at vertex *v* (*v* = 1, … *V*) of a hemisphere of the brain from subject *j* (*j* = 1, …, *n*_*i*_) in scanner *i* (*i* = 1, …, *M*). Let **x**_*ij*_ = (*x*_*ij*1_, …, *x*_*ijq*_)^*T*^ be the *q*-dimensional covariate vector for subject *j* in scanner *i* (e.g., age and sex). We model *y*_*ijv*_ by

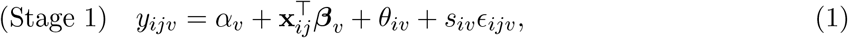

where *α*_*v*_ is the intercept, ***β***_*v*_ is the regression coefficient vector, *θ*_*iv*_ is the scanner-specific intercept for scanner *i*, 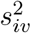 is the scanner-specific variance for scanner *i*, and *ϵ*_*ijv*_ is the noise term with the unit variance. Note that, at the vertex level, the Stage 1 model is equivalent to ComBat’s specification of batch effects (Johnson et al., 2007; Fortin et al., 2018) that models heterogeneous means and variances across scanners.

#### 2.1.2 Characterization of heterogeneous spatial covariances

To account for spatial dependence of cortical thickness, we model ***ϵ***_*ij*_ = (*ϵ*_*ij*1_, …, *ϵ*_*ijV*_)^*T*^ using the spatial Gaussian process (GP), that is, 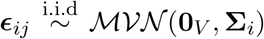 for *j* = 1,...,*n*_*i*_. Specifically, we decompose ***ϵ***_*ij*_ into three additive random effects as

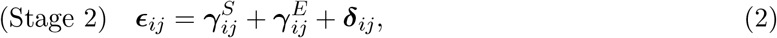

with an assumption that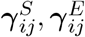, ***δ***_*ij*_ are independent to each other.

- 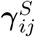 and 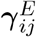 are spatial random effects that are modeled by 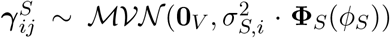 and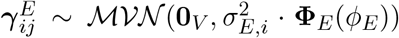 Here, we assume that the covariances of 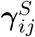and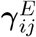 are characterized by (i) heterogeneous spatial variances which represented by scanner-specific parameters 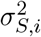 and 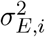 and (ii) homogeneous underlying autocorrelations across scanners which represented by the spatial autocorrelation functions (SACFs). For vertices *v* and *v*^*∗*^,

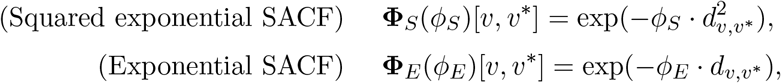

where *d*_*v,v*_*∗* is the geodesic distance between vertices *v* and *v*^*∗*^, and the scanner-invariant parameters *ϕ*_*S*_ and *ϕ*_*E*_ determine how fast the spatial autocorrelation decreases with distance. The exponential SACF falls off rapidly with small distances but then tails off much slower than the squared exponential SACF as distance increases. Combining these two forms provides the flexibility to simultaneously account for spatial correlation at shorter distances through the squared exponential SACF and capture the heavy-tailed nature of spatial dependence within the brain through the exponential SACF. The integration of squared exponential and exponential SACFs has shown its utility when modeling spatial autocorrelations in fMRI data (Cox et al., 2017).
- ***δ***_*ij*_ is the non-spatial effect modeled by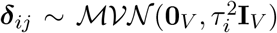. Its covariance structure 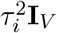 includes scanner-specific parameter 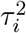 represents heterogeneous non-spatial variances across scanners.

Altogether, the marginal covariance of 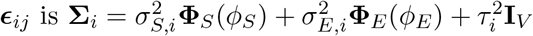

### 2.2 Spatial autocorrelation normalization (SAN)

#### 2.2.1 Stage 1

The Stage 1 model requires estimating *α*_*v*_, ***β***_*v*_, *θ*_*iv*_ and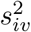, with an important data-specific consideration that the additive and multiplicative batch effects (*θ*_*iv*_ and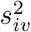) would be smooth over space. Considering such smoothness in the estimation step would reduce the variance of the parameter estimates, which is expected to improve the performance of the Stage 2 of SAN especially when the number of subjects is small.

To achieve this, SAN pools information within a series of prespecified local neighbors (Wang et al., 2021). Specifically, we define a local neighbor 𝒩_*r*_(*v*) for each vertex as the set of vertices whose geodesic distances from vertex *v* are less than or equal to *r*, and apply ComBat (Johnson et al., 2007; Fortin et al., 2018). In ComBat, we obtain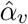, 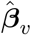 and residuals *ê*_*ijv*_ via least squares, and then impose normal-inverse-gamma priors to *θ*_*iv*_ and 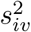 and estimate them using the empirical Bayes approach, which shrink these estimates towards the overall means and variances of 𝒩_*r*_(*v*).

The performance of ComBat depends on the selection of the radius *r*. When *r* = 0mm, we estimate *θ*_*iv*_ and 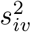 separately for each vertex without borrowing any information from local neighbors.

When *r* = ∞, we estimate *θ*_*iv*_ and 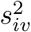 by using all vertices in the data, which would result in ‘too smooth’ estimates shrunk towards the brain-level means and variances. An appropriate level of *r* needs to be chosen by considering the bias-variance tradeoff (Wang et al., 2021). In this paper, we choose ComBat model with *r* = 5mm as a default to borrow information across neighbors without inducing too much bias. A justification of this choice is provided in the supplementary material S1.

#### 2.2.2 Stage 2

We generalize the covariance regression proposed by Zou et al. (2017) to estimate 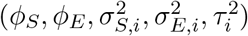.

The covariance regression approach uses the method of moments (MoM) estimators, which provide a highly computationally efficient and consistent estimator of the variance component parameters, with some loss in efficiency, compared to the likelihood-based methods. We note that the likelihood-based methods would be intractable as the number of vertices gets larger or the number of scanners/sites increases (Zou et al., 2017). Previous work from Park & Fiecas (2022) has also shown empirically that the MoM estimators yielded nearly unbiased estimates of parameters. In SAN, the parameters are estimated by minimizing the following objective function:

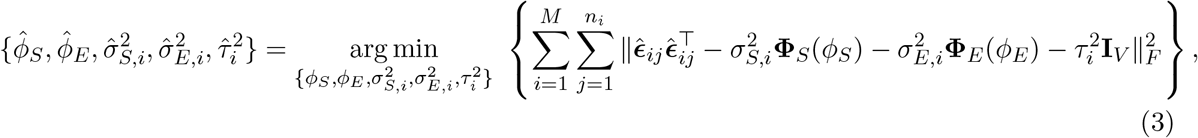

where 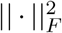 is the squared Frobenius norm of a matrix. For optimization, we first note that, when *ϕ*_*S*_ and *ϕ*_*E*_ are given (so that **Φ**_*S*_ *≡* **Φ**_*S*_(*ϕ*_*S*_) and **Φ**_*E*_ *≡* **Φ**_*E*_(*ϕ*_*E*_)), a closed-form solution for other parameters minimizing the objective (3) is provided by

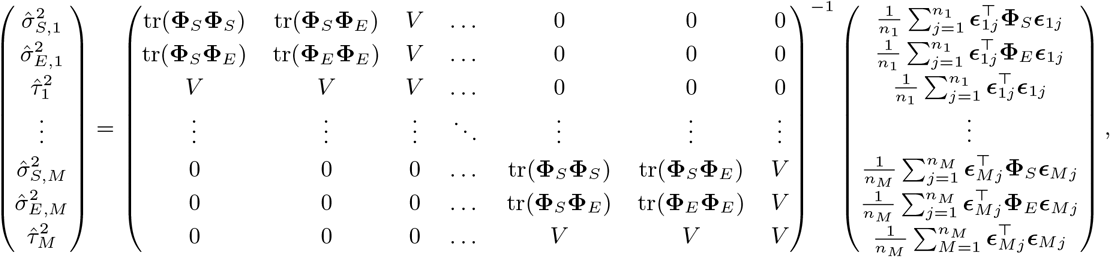

where tr(*·*) is the trace of a matrix. Subsequently, when we plug in 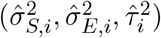 to the closedform solution in Equation (3), it simplifies the optimization process, reducing it to finding the minimum of the updated objective function with respect to (*ϕ*_*S*_, *ϕ*_*E*_). This step reduces the number of parameters for the optimization from 2 + 3*M* to 2, significantly enhancing the computational efficiency regardless of the number of scanners (*M*) used in a study. In our implementation in R, the Nelder-Mead method (Nelder & Mead, 1965) was used to solve this nonlinear optimization problem.

Under the GP assumption, the conditional expectations of 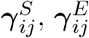 and ***δ***_*ij*_ are given by

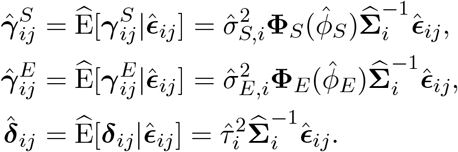

#### 2.2.3 Construction of harmonized data

With all decompositions and estimations made in both stages, we remove scanner-specific means and normalize scanner-specific covariances to construct the harmonized data. Since the sources of covariance heterogeneity are characterized by scanner-specific parameters 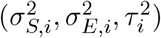, we normalize 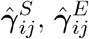 and 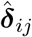 by

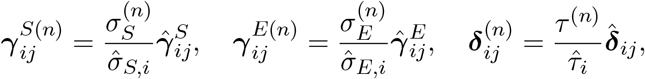

where 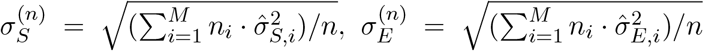 and 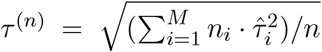.

Therefore, the final normalized data is given by

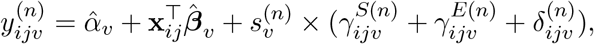

where 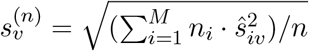.

### 2.3 Integrating SAN with other harmonization methods

While the primary focus of SAN lies in modelling and normalizing scanner-specific spatial autocorrelations, it is worth noting that spatial heterogeneity might not be the only source of the scanner effects. In such a case, SAN’s formulation could suffer from oversimplification, failing to address the full complexity associated with heterogeneous non-spatial variations. Therefore, we considered applying covariance harmonization methods to 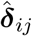 from Stage 2 of SAN to capture potential remaining scanner effects. In this paper, we consider CovBat (A. A. Chen et al., 2022) (“SAN+CovBat”) and RELIEF (Zhang et al., 2023) (“SAN+RELIEF”) as these are methods primarily developed to harmonize covariances. In our data analysis, we did not find any noticeable differences in performance between SAN and SAN+CovBat or SAN+RELIEF. Detailed analysis results of SAN+CovBat and SAN+RELIEF are provided in the supplementary material S3.

## 3 Data Analysis

### 3.1 Data preparation and preprocessing

We used cortical thickness data from the Social Processes Initiative in the Neurobiology of the Schizophrenia(s) (SPINS) study to evaluate SAN’s performance. The SPINS study includes multi-modal neuroimaging data in individuals diagnosed with schizophrenia spectrum disorders (SSDs) and control participants. Participants aged 18-59 were recruited for SPINS between 2014-2020. All participants signed an informed consent agreement, and the protocol was approved by the respective research ethics and institutional review boards. All research was conducted in accordance with the Declaration of Helsinki. See Viviano et al. (2018) and Oliver et al. (2021) for details.

Scans were obtained from three different imaging sites: Centre for Addiction and Mental Health (CAMH), Maryland Psychiatric Research Center (MPRC), and Zucker Hillside Hospital (ZHH). General Electric (GE) 3T MRI scanners were used at CAMH and ZHH (750w Discovery and Signa, respectively), while MPRC used the Siemens Tim Trio 3T (ST). However, in the third year of the study, all sites switched to the Siemens Prisma (SP) scanner. Given the limited number of samples from the Siemens Tim Trio 3T (60 samples), we excluded ST data and focused our analysis on the two scanner types: GE and SP. Our final dataset comprised 357 subjects across these two scanner types: 164 were imaged on the GE (66 females, 104 patients, aged 18-55) and 193 on the SP (79 females, 105 patients, aged 18-55) after visual quality control.

MRI data preprocessing was performed using fMRIPrep 1.5.8 (Esteban et al., 2019), based on Nipype 1.4.1 (Gorgolewski et al., 2011). T1-weighted images were corrected for intensity non-uniformity and skull-stripped using ANTs (Avants et al., 2008). Cortical surfaces were reconstructed using FreeSurfer 6.0.1 (Dale et al., 1999). Cortical thickness data were resampled to fsaverage5 space, including Gaussian smoothing with FWHM of 0, 5, and 10 mm, leaving 9,354 cortical vertices for the left hemisphere and 9,361 for the right hemisphere after excluding vertices on the medial wall. Upon visual inspection of the preprocessed T1 images, we excluded 29 images that did not pass quality control or passed with small issues, including minor bad skull stripping, MNI warping, and tiny under-inclusive Freesurfer masking. The pairwise geodesic distance was computed from the pial surface to reflect surface geometry.

### 3.2 Evaluations of covariance heterogeneity across scanners

Several metrics have been proposed to compare harmonization methods (Hu et al., 2023). These metrics typically test inter-scanner differences in means or variances (e.g., density estimations, MANOVA, t-tests), visualize low-dimensional differences (e.g., PCA), or indirectly evaluate predictive performances using machine learning methods (e.g., random forest). However, we note that (i) none of these explicitly assess inter-scanner *covariance* heterogeneity, and (ii) these are mostly limited to the low-dimensional features. Therefore, we introduce two new metrics which are “direct” quantification of differences in covariances that can be easily visualized and statistically explained.

#### Covariance Analysis of Scanner Heterogeneity (CASH score)

The CASH score, inspired by the SWISS score (Cabanski et al., 2010), extends the idea of Analysis of Variance (ANOVA) in addressing covariance differences across scanners from multiple features. By treating empirical covariance measurements as observations in the ANOVA framework, CASH score computes the Total Sum of Squares (SST) as the sum of squared differences between each covariance measurement and the overall mean of the covariance measurement across scanners. SST can be further decomposed into Inter-scanner Sum of Squares (ISS) and Within-scanner Sum of Squares (WSS), where ISS quantifies the difference in covariance measurements between scanners and WSS captures the variability within each scanner. CASH score quantifies the relative contributions of ISS and WSS in empirical covariance measurements and adjust for degrees of freedom. A higher CASH score indicates severe inter-scanner biases in covariances. When 𝒩 is the set of pairs of features, the formula for the CASH score is

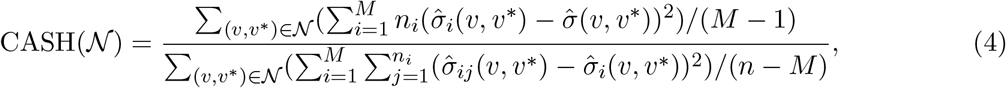

Where 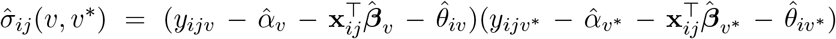. Here, 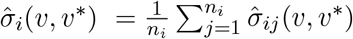 represents the covariance measurement obtained from scanner *i* between vertex *v* and *v*^*∗*^, and 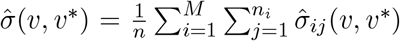 is the pooled covariance measurement across all scanners between vertex *v* and *v*^*∗*^.

#### Covariogram ratio

Covariogram quantififes how the covariance of spatial features depends on the distance (Banerjee et al., 2003). Specifically, for each image, the covariogram for a distance *d* is defined as:

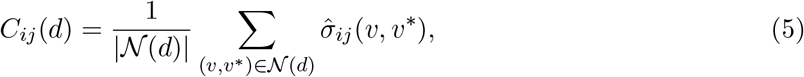

where 𝒩 (*d*) is the set of vertex pairs such that their distance is between *d − η* and *d* + *η* for a small bandwidth *η >* 0. | *𝒩* (*d*)| is the number of elements in this set.

We define the covariogram ratio as the ratio of covariograms computed from harmonized data for each scanner to those obtained from the original, pooled data across all scanners. We first obtain the pooled average of empirical covariagrams, provided by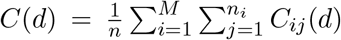, and the scanner-specific average by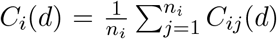 For harmonized data, we obtain the formula is provided by scanner-specific average by 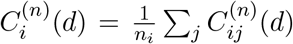by replacing *y*_*ijv*_ with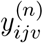. From here, the formula is provided by

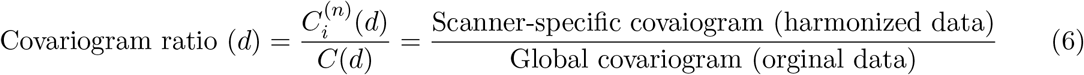

The ratio closer to 1 indicates better preservation of underlying homogeneous spatial variations, as the scanner-specific covariogram shrinks towards the global covariogram after harmonization.

### 3.3 Smoothing increases heterogeneity in covariances across scanners

In terms of cortical thickness data preprocessing pipelines, there is currently no consensus on whether smoothing should be done before or after harmonization. This study provides empirical evidence that spatial smoothing can exacerbate the complexity of covariance heterogeneity among different scanners. For the current context, we use CASH score and define 𝒩= 𝒩_*r*_(*v*) with *r* = 5mm, denoting the local neighbors for a vertex *v*.

We use three sets of raw cortical thickness data (after regressing out scanner-specific means): unsmoothed, 5mm smoothed, and 10mm smoothed. From Figure 1, we first observe from results of the unsmoothed data that covariance heterogeneity is localized and prominent in regions including, but not limited to, pericalcarine, caudal anterior cingulate, paracentral, precentral, postcentral, superior temporal, midtemporal, and insula and entorhinal cortices. We also see that smoothing not only intensifies inter-scanner covariance biases but also spreads these biases to a larger spatial extent compared to unsmoothed data. This effect is further magnified when a larger smoothing kernel is used. We illustrate in a simplified setting with mathematical proof in the supplementary material S2 to show how spatial smoothing can amplify the covariance heterogeneity between different scanners. Therefore, we used the unsmoothed data to evaluate SAN’s performance throughout this paper, with sensitivity analysis regarding the effect of smoothing provided in Section 3.7.

**Figure 1:**
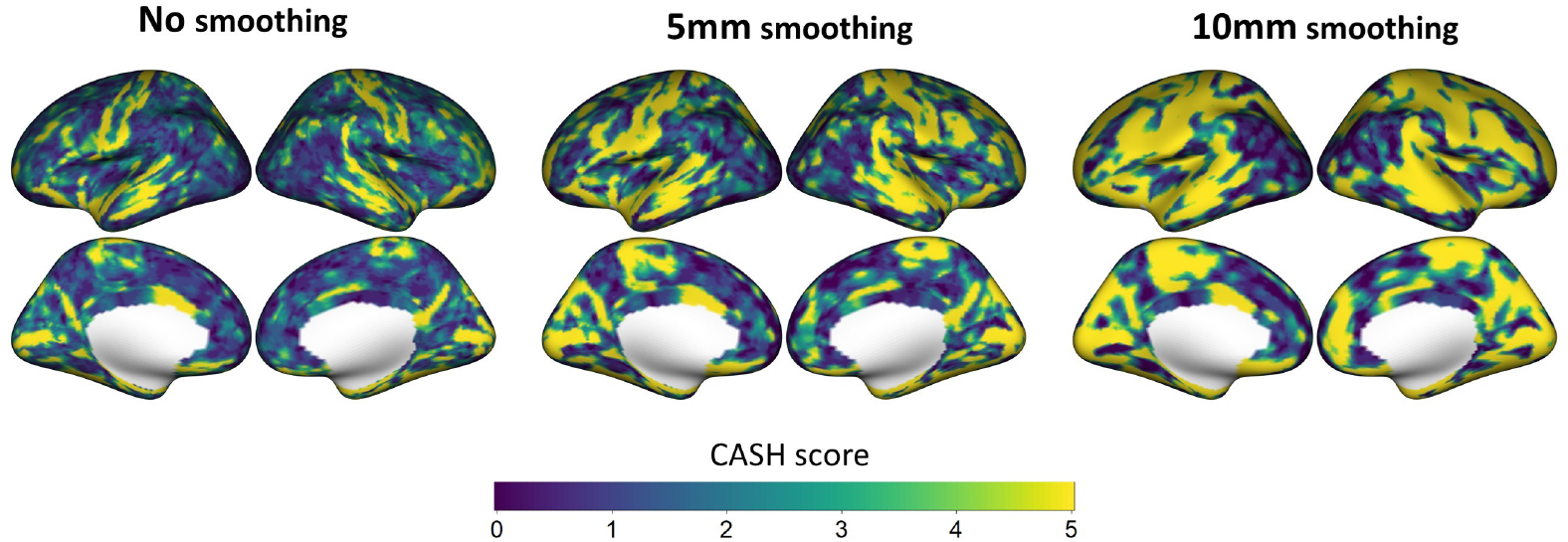
CASH score brain maps obtained from unsmoothed and smoothed data before applying any harmonization methods. A larger CASH score at a vertex implies higher covariance heterogeneity in local neighbors surrounding the vertex. The color bar was capped at 5 for better visualizations.

### 3.4 Brain-level analysis

#### 3.4.1 Covariogram analysis

In Figure 2(a), when considering the covariogram ratios for both GE and SP within the range of 0mm to 40mm, SAN consistently outperforms the other competitors across most distances. While CovBat effectively aligns the SP-averaged covariogram closer to the pooled covariogram, it introduces more deviations in GE when distances exceed 20mm. Raw and ComBat harmonized data exhibit similar covariogram ratios, consistently larger than those of SAN. Notably, RELIEF displays distinct behavior compared to the other methods; it demonstrates the smallest ratios for SP but the largest ratios for GE, even surpassing raw data in the latter case. This observation suggests that RELIEF’s performance exhibits an imbalance across scanners, potentially arising from an excessive elimination of spatial variation specific to GE.

**Figure 2:**
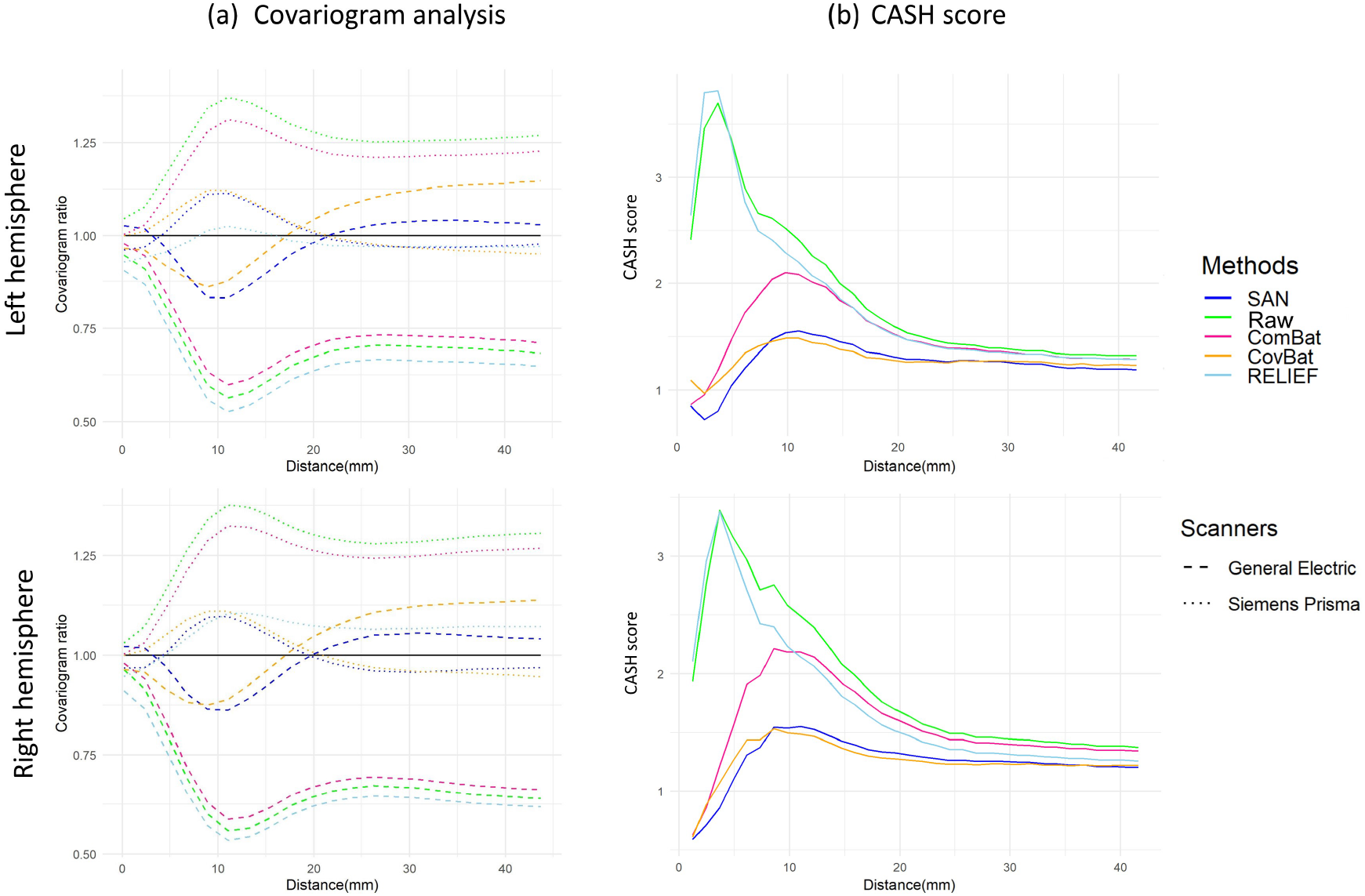
Summary of covariogram ratios and CASH scores obtained from different harmonization methods. The left panel shows covariogram ratios, with the solid horizontal line representing a ratio of 1. The right panel shows CASH scores.

#### 3.4.2 CASH score analysis

We evaluate how CASH scores depend on distances. Specifically, we replace 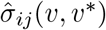 and 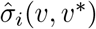 with 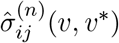 and 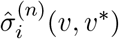 for each harmonized data, and use *𝒩*= *𝒩* (*d*) as provided in Section 3.4.1. This evaluates how the average degree of covariance heterogeneity in the brain depends on distances. As shown in Figure 2(b), noticeable performance differences among harmonization methods are observed within the 0-20mm range. Specifically, SAN and CovBat exhibit markedly smaller CASH scores compared to other methods. Within this range, our methods outperform CovBat within the 10mm interval. However, for distances ranging from 10mm to 20mm, CovBat shows slightly smaller CASH scores. Beyond the 20mm threshold, all methods demonstrate similar performance, with decreasing CASH scores. It could be because (spatial) heterogeneity in covariances would be marginal as the distance increases. Still, SAN consistently produce slightly smaller CASH scores, supporting its empirical performance.

#### 3.4.3 Brain maps of CASH scores

To visualize the decrease in covariance heterogeneity between scanners both globally and locally on the brain surface after harmonization, we compute CASH scores for each harmonization method as done in Section 3.3. Figure 3 shows the brain maps of CASH scores. Overall, SAN shows great improvement in reducing CASH scores compared to other methods. Covariance heterogeneity between scanners remains prominent in RELIEF brain maps. ComBat and CovBat perform slightly better, though some minor spatial clusters still present large CASH scores.

**Figure 3:**
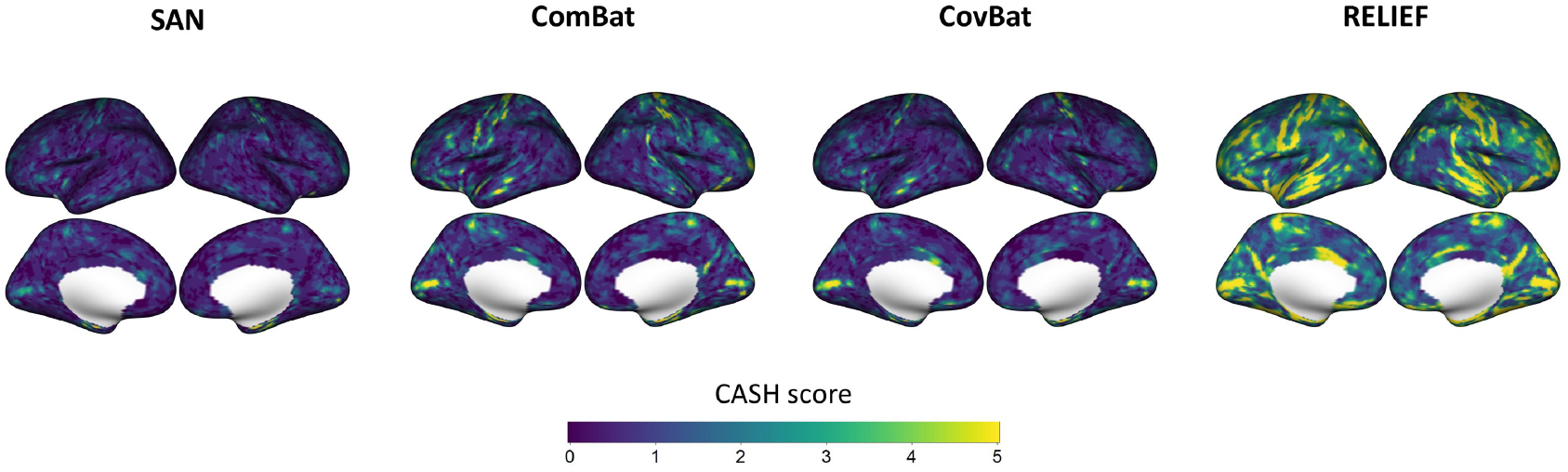
Brain maps of CASH scores obtained from different harmonization methods. The values exceeding 5 are capped for visualizations.

#### 3.4.4 Retaining biological variability

To evaluate how well SAN preserves biological variation in the cortical thickness data, we first select a subset of vertices that exceed the 80th percentile of the CASH scores, which result in 1871 and 1873 vertices from the left and right hemispheres, respectively. Then, for each harmonized data, we use GLM for each feature by using 4 covariates (age, sex, diagnosis, age *×* diagnosis) and obtain *p* values. Then we compute the proportion of the significant vertices associated each covariate (FDR-adjusted *p* values *≤* 0.05).

For covariates other than age, no significant associations are observed in either hemisphere across all harmonization methods. For age, the proportion of the significant vertices in the left hemisphere is 0.86% (SAN), 0.91% (Raw), 0.86% (ComBat), 1.02% (CovBat) and 0.96% (RELIEF). In the right hemisphere, it is 1.98% (SAN), 1.98% (Raw), 1.98% (ComBat), 1.60% (CovBat) and 2.46% (RELIEF). The proportion of the significant vertices in all harmonized data appeared to be indifferent, which supports that SAN retains similar levels of biological information to other harmonization methods.

### 3.5 Within-ROI analysis

The purpose of this section is to investigate how brain-wise spatial harmonization affects locally within a predefined region. We use Desikan-Killiany Atlas (Desikan et al., 2006), which includes 34 regions of interest (ROIs) in each hemisphere (after excluding corpus callosum). With each harmonized dataset, we segment the cortical thickness values into 34 subsets for the left hemisphere according to this atlas.

Figure 4 shows the barplots of CASH scores for the top 10 regions in each hemisphere with the highest CASH scores from the raw data. Overall, these regions are highly overlapped between two hemispheres and relatively smaller in size, having 142 vertices for the left hemisphere and 173 vertices for the right hemisphere on average, which is less than the average of 265 vertices for all regions. This observation is also consistent with our findings in section 3.4, where the inter-scanner heterogeneity in covariances is most pronounced at shorter distances. Also, among these regions, SAN shows the lowest CASH scores for 6 regions in the left hemisphere and for 5 regions in the right hemisphere. CovBat showed the lowest CASH scores for 3 regions in the left hemisphere and 5 regions in the right hemisphere, which partially supports that there might be some degrees of spatial nonstationarity in cortical thickness data such that SAN (which assumes spatial stationarity) would perform worse than CovBat in some localized areas. Lastly, the performance of ComBat was poor in harmonizing covariances within most ROIs. While ComBat shows the lowest CASH score for caudal anterior cingulate, the disparities between ComBat and SAN are marginal.

**Figure 4:**
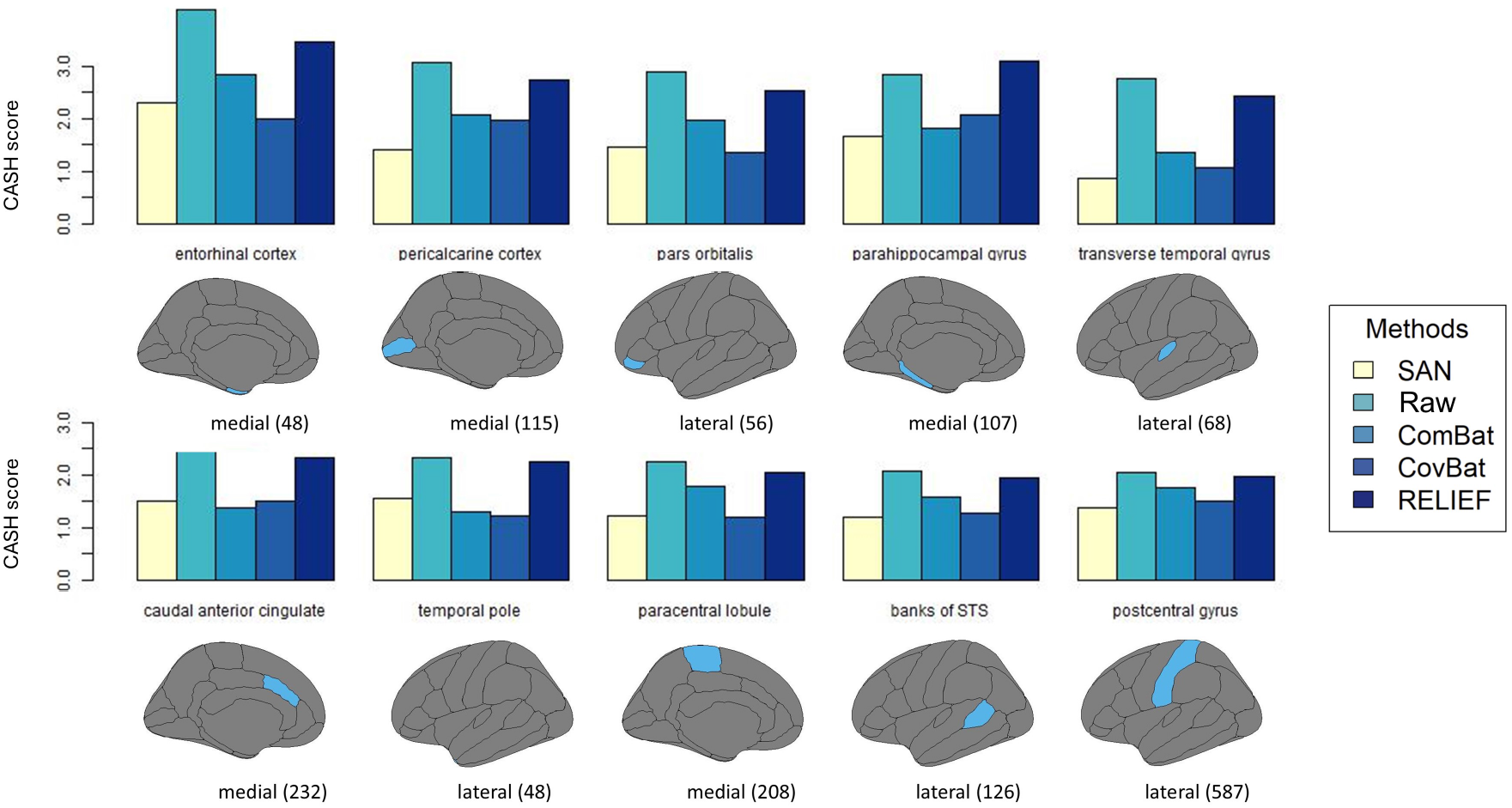
ROI-level CASH score results. The top 10 regions in the left hemisphere are selected based on their highest CASH scores from the raw data. For each region, the CASH score is computed using Equation 4, using all pairs of vertices within that region as the vertex set *N*. Below the corresponding bar plots, diagrams of the ROI complete with vertex counts are displayed. The results for the right hemisphere are provided in the supplementary material S5.

### 3.6 Between-ROI analysis

We investigate whether brain-wise harmonization before parcellation is effective in homogenizing covariances in region-level data. Using Desikan-Killiany atlas, we derive ROI-level cortical thickness data by averaging cortical thickness values within each corresponding region. In this section, we assess two types of harmonized data: (i) the vertex-level harmonization followed by parcellation and (ii) partitioning the data to construct the average cortical thickness for each ROI and then harmonizing the ROI-level cortical thickness data. For (ii), we use ComBat, CovBat, and RELIEF (denoted as ROI-level ComBat, ROI-level CovBat and ROI-level RELIEF). Although vertex-level data may not be directly applicable to ROI-level analyses, the advantages observed in capturing and retaining the underlying shared spatial correlation across scanners can still be leveraged when working with ROI-level data by representing averaged measurements within regions.

In addition to the CASH score, we also use QDA to assess the predictive capability of each harmonized data in relation to scanners. It is important to note that a harmonization method that exhibits superior performance in mitigating scanner effects will yield inferior predictive performance. Similarly to Zhang et al. (2023), we choose QDA because it relies solely on mean vectors and covariance matrices. Therefore, any variations in predictive performance can be directly attributed to the harmonization of scanner-specific means and covariances. Using leave-one-out cross-validation, we calculate the average accuracy as well as ROC curve’s area under the curve (AUC) for each harmonized dataset after regressing out covariate effects.

The results for the CASH scores, accuracy and AUC for scanner prediction are shown in Table 1. For CASH scores, the ROI-level ComBat and ROI-level CovBat exhibit the most effective performance in minimizing inter-scanner variabilities. It can be attributed to the favorable utilization of ROI-level data in ROI-level ComBat/CovBat for harmonization. However, SAN shows noticeably smaller CASH scores than all other vertex-level harmonization methods. This suggests that our method excels in recovering underlying homogeneous spatial correlations across scanners. For evaluating the predictive performance of scanners, we see that ROI-level RELIEF and vertex-level RELIEF achieve the lowest prediction accuracy and AUC values, which is consistent with the findings by Zhang et al. (2023). Considering that RELIEF performed worse in reducing the CASH scores, this result suggests that an optimal harmonization method should be chosen carefully based on the purpose of the desired research (e.g., prediction vs. inference).

**Table 1:** The summary of CASH score, the accuracy and AUC of predicting scanners. The smallest values for vertex-level/ROI-level methods are bolded.

| Measure | CASH score | Accuracy | AUC |
| --- | --- | --- | --- |
| SAN | **2.351** | 0.608 | 0.601 |
| Raw | 4.546 | 0.641 | 0.628 |
| ComBat | 3.648 | 0.605 | 0.593 |
| CovBat | 4.152 | 0.622 | 0.61 |
| RELIEF | 4.349 | **0.529** | **0.517** |
| ROI-level ComBat | **1.746** | 0.605 | 0.602 |
| ROI-level CovBat | 1.823 | 0.555 | 0.549 |
| ROI-level RELIEF | 3.827 | **0.527** | **0.517** |

### 3.7 Sensitivity analysis of the impact of smoothing

To understand the impact of smoothing on harmonization performance, we investigate the sensitivity of harmonization methods to varying levels of smoothing.

#### 3.7.1 Smooth data first then harmonize

We use the same 5mm smoothed data and 10mm smoothed data as Section 3.3 and apply different harmonization methods. Figure 5(a) shows CASH score brain maps derived from data smoothed prior to harmonization. Brain maps corresponding to the 5mm smoothing level are similar to those observed in unsmoothed harmonized data. SAN outperforms other harmonization methods in reducing inter-scanner variabilities. However, it performs poorly with the 10mm smoothing, as indicated by the appearance of large spatial clusters characterized by notably high CASH scores on the maps. This limitation is likely due to the nonstationarity induced by spatial smoothing, which could lead to a severe violation of the stationary Gaussian process assumption made by SAN. Using a FWHM=10mm may introduce or exaggerate nonstationarity at broader scales. It is also worth noting that ComBat (which does not address covariance heterogeneity) seems to be the best model for harmonizing covariances when 10mm smoothing is applied. Therefore, we note that SAN is preferably applied to unsmoothed data or data with minimal smoothing levels (e.g., less than 5mm) to ensure optimal harmonization outcomes.

**Figure 5:**
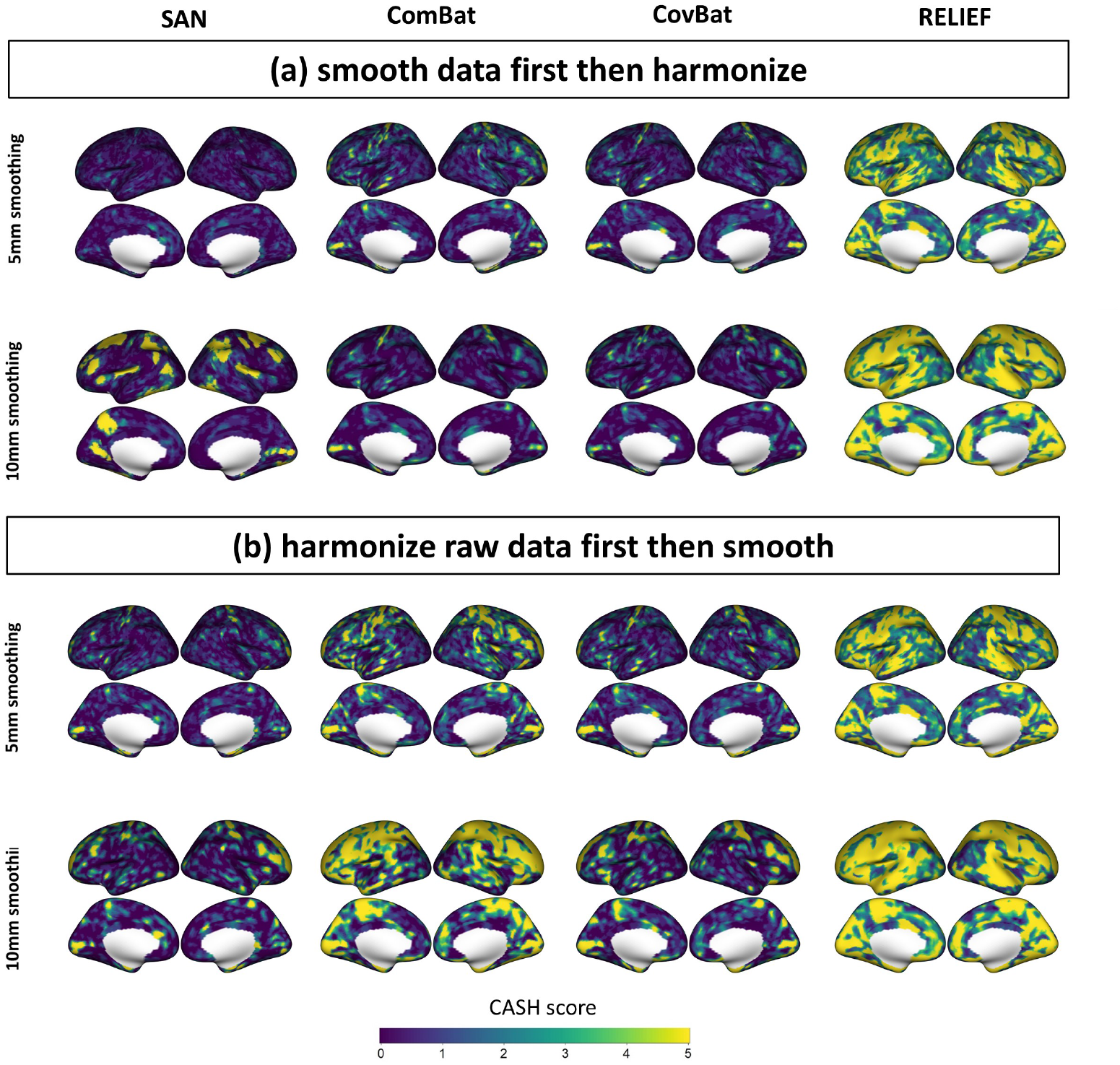
CASH scores computed from data smoothed prior to harmonization (Figure (a)) data harmonized prior to smoothing (Figure (b)).

#### 3.7.2 Harmonize raw data first then smooth

We apply Gaussian smoothing with FWHM of 5 and 10mm after exporting the data to cifti format. The CASH score brain maps, derived from data harmonized prior to smoothing, are shown in Figure 5 (b). As shown in Section 3.3, the increase in smoothing levels leads to larger inter-scanner covariance biases, both in terms of magnitude and spatial extent, compared to unsmoothed data. Compared to unsmoothed data in Figure 3, more prominent inter-scanner covariance biases are present to a larger spatial extent in Figure 5 (b). Given that no harmonization method is capable of completely eliminating inter-scanner covariance biases, smoothing may exacerbate and propagate these residual biases. However, it is worth noting that SAN shows the least susceptibility to the negative influence of post-smoothing on harmonization compared to other methods that neither address covariance heterogeneity nor account for spatial covariances.

### 3.8 Data-driven simulations

To adequately assess the precision and uncertainty of the harmonization method, we conduct a data-driven simulation using cortical thickness data from the SPINS study. Data-driven simulations are preferred in scenarios where the complex nature of target data makes it challenging for parametric simulations to replicate, as evidenced by previous neuroimaging method validations (Park & Fiecas, 2022; Weinstein et al., 2022; Pan et al., 2024). In our case, SAN assumes that a parametric, isotropic, stationary Gaussian process perfectly captures covariance heterogeneity between scanners, which could be an oversimplified representation of real cortical thickness data. Conducting a parametric simulation based on SAN’s assumption could bias results in favor of SAN while failing to capture the true characteristics of cortical thickness data. To address these limitations and ensure a fair evaluation of SAN’s performance even under potential misspecification (e.g., non-stationarity), we conduct a data-driven simulation that enables us to generate simulated datasets that closely resemble real cortical thickness data, thereby capturing underlying complexities more effectively.

We repeat the following procedure 1,000 times, generating a total of 1,000 simulated datasets. Each simulated data involves randomly selecting 75 individuals from GE and 75 individuals from SP. To streamline computations and enhance result presentation, we introduce some modifications in the calculation of CASH scores. Given the large number of simulation samples, we select distance ranges as: 0mm-5mm, 5mm-10mm, 10mm-15mm, 15mm-20mm, 20mm-30mm and 30mm-40mm. With each of these ranges, we compute this measure using vertex pairs that fall within the specified range. For 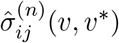 and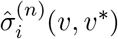, we apply harmonization methods to each simulation dataset and then calculate scores as done in section 3.4.2. For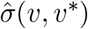, we use the overall original dataset to estimate the underlying pooled covariances.

Figure 6 shows boxplots of CASH scores from bootstrapping, which align with the findings in Figure 2(b). Across the six ranges, SAN consistently exhibited smaller medians of CASH scores compared to other methods, particularly for CASH scores below 15mm. This can be attributed to its ability to capture and harmonize most of the scanner-specific local spatial dependencies in the data. Following it are CovBat and RELIEF. CovBat shows comparable performance to SAN in the 15mm-30mm range, while RELIEF shows similar CASH scores to SAN in 30mm-40mm.

**Figure 6:**
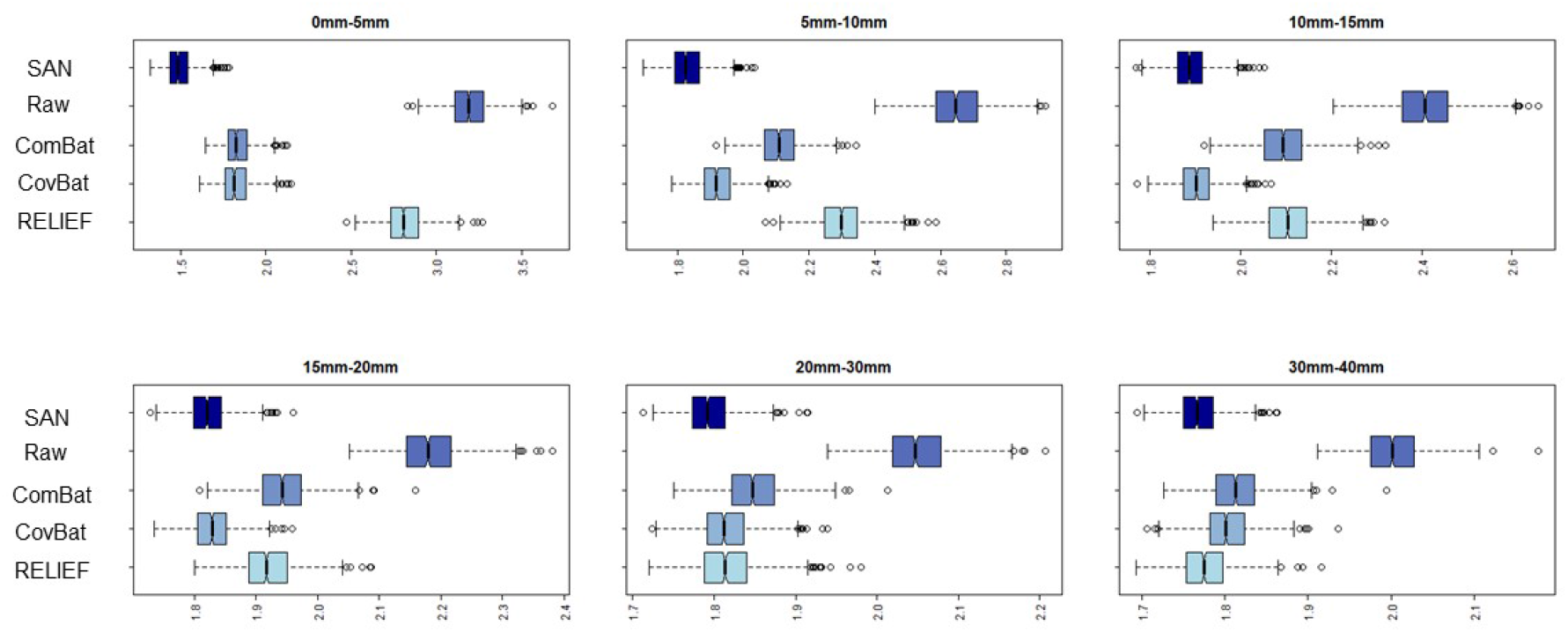
Boxplots illustrating CASH scores of raw data and different harmonized data across six distance intervals, derived from 1000 simulated results. SAN shows smaller medians of CASH scores than other methods across intervals.

## 4 Discussion

In this paper, we propose a new harmonization method, SAN, that identifies and parameterizes sources of heterogeneity in vertex-level cortical thickness data collected from different scanners or sites. We use Gaussian process to model and homogenize spatial covariances, and its probabilistic modeling ensures the smoothness of the harmonized cortical thickness data, which existing methods do not provide. SAN aligns with the growing need for providing high-quality data to downstream whole-brain analysis, especially those involving spatial covariance modelling. Despite several “covariance harmonization” methods proposed in the neuroimaging literature to date, our analysis of the SPINS study highlights that the performance of these other methods is highly variable and often suboptimal for vertex-level cortical thickness data, which can be attributed to the absence of leveraging spatial information to capture covariance heterogeneity. The promising performance of SAN on vertex-level cortical thickness data supports the need to develop statistical harmonization methods tailored to specific neuroimaging data types (e.g., network structures in functional connectivity), revealing their specific covariance patterns.

Our data analysis with CASH scores suggests that there are specific anatomical regions that reveal a high degree of covariance heterogeneity across scanners, for which SAN’s brain-level harmonization successfully homogenized without compromising biological variability inherent in the data. Although initially designed for vertex-wise data, SAN also proves advantageous for within-ROI and between-ROI analyses by ensuring scanner-specific spatial normalization at the foundational level, which is critical for constructing accurate ROI-level metrics. The data-driven simulation further supports the effectiveness and robustness of SAN.

The sensitivity analysis regarding the impact of smoothing on harmonization performance underscores the robustness of SAN against preprocessing choices. While smoothing is a common preprocessing step in neuroimaging analysis, we provide empirical and theoretical evidence of higher mean and covariance heterogeneity induced by smoothing in multi-site/scanner studies. Our findings suggest that smoothing before harmonization causes covariance harmonization methods (SAN, CovBat, RELIEF) less effective, with ComBat performing better on highly smoothed data. Harmonizing before smoothing, on the other harnd, exacerbates inter-scanner covariance biases to harmonized data, yet SAN remains the least impacted by subsequent smoothing. SAN offers flexibility in its order of application before or after smoothing. For minimal smoothing levels (e.g., less than 5mm FWHM), SAN can be applied after smoothing, whereas for higher levels of smoothing (e.g., 10mm FWHM), SAN can be applied before smoothing without inducing larger inter-scanner biases observed in other harmonized data. In terms of the optimal data processing pipeline, more empirical and theoretical evidences are needed.

SAN’s harmonization aims to recover an optimal ‘pooled’ covariance from the parametrizations of the Gaussian process, which reduces the CASH score significantly than other harmonization methods we considered. However, a cautionary note is needed in choosing the harmonization method that meets users’ research purposes. SAN is recommended when covariance modeling of the vertex-level cortical thickness data is critical in improving statistical inference. In contrast, RELIEF performs best in impeding the detection of scanners at the ROI level. However, RELIEF does not seem effective in reducing the CASH score as RELIEF primarily focuses on *removing* scanner-specific latent factors. These results suggest that, despite higher power shown in Zhang et al. (2023) in massive univariate analysis, SAN would perform more promisingly in the spatial-extent inference that requires explicit spatial aucorrelation modeling.

SAN has room for improvement. First, we downsampled cortical thickness data to fsaverage5 space (*V ≈* 10, 000) in our analysis, which seems to be sufficient in capturing dense spatial information without significant loss of information (Mejia et al., 2019; Xu et al., 2016). However, SAN would be computationally limited when applied to higher resolutions (e.g., fsaverage6 or fsaverage7). Although implementing SAN is computationally feasible when *V ≈* 10, 000, more research is needed to make it even more computationally efficient in higher dimensions. Second, longitudinal studies (e.g., Alzheimer’s Disease Neuroimaging Initiatives) have identified inter-scanner biases in cortical thickness data (Beer et al., 2020). Extending SAN to longitudinal neuroimaging studies would simultaneously account for both the within-subject variability and covariance heterogeneity across scanners. This enhancement could maintain within-subject dependencies in the harmonization process, thereby improving data quality for longitudinal designs not currently addressed by our existing SAN framework. Finally, there are also other structural imaging metrics based on the human cerebral cortex, such as surface area, and gyrification, which have shown discrepancies in their measurements across scanners (Iannopollo et al., 2021; de Moraes et al., 2022; Radua et al., 2020). Although this paper primarily focuses on harmonizing cortical thickness data, future research could explore the validity of SAN in these imaging modalities.

## 5 Software

The R package for implementing SAN is publicly available at https://github.com/junjypark/SAN. Our harmonization took approximately 1 hour on a Macbook Air (M2,2022) with 16GB RAM to harmonize data with 9,354 imaging features from 357 subjects, which supports the computational feasibility of the proposed method. For server users, the harmonization was completed in approximately 36 minutes on a server cluster node (Lenovo SD350) equipped with 40 Intel “Skylake” cores (2.4GHz) and 202GB RAM. Parallel computing is supported by the package to mitigate computational costs working with a large number of scanners or sites.

## Data and Code Availability statement

The R package for implementing SAN is publicly available at https://github.com/junjypark/SAN.

## Acknowledgements

The SPINS study was supported by the National Institute of Mental Health (1/3R01MH102324-01, 2/3R01MH102313-01, 3/3R01MH102318-01). RZ was supported by the Doctoral Fellowship from the University of Toronto Data Science Institute. LDO was supported by the Brain & Behavior Research Foundation. ANV was supported by the National Institute of Mental Health (1/3R01MH102324 & 1/5R01MH114970), Canadian Institutes of Health Research, Canada Foundation for Innovation, CAMH Foundation, and University of Toronto. JYP was supported by Natural Sciences and Engineering Research Council of Canada (NSERC) (RGPIN-2022-04831), the University of Toronto’s Data Science Institute (Catalyst Grant), McLaughlin Centre (Accelerator Grant), and the Connaught fund. The computing resources were enabled in part by support provided by University of Toronto and the Digital Research Alliance of Canada (alliancecan.ca).

