## Supplementary file for "SAN: mitigating spatial covariance heterogeneity in cortical thickness data collected from multiple scanners or sites"

### S1 The bootstrapping procedure for deriving $r$ for the Stage 1 of SAN

We conduct bootstrap to select an optimal radius $r$ from a candidate set 0mm, 5mm, 10mm, and 20mm. The elements in the set can be specified by the investigators a priori. For each bootstrap $b$, the procedure is summarized by (i) randomly selecting 75 subjects from each scanner type respectively; (ii) fitting ComBat model with bootstrap data, and obtaining scanner-specific means $\hat{\theta}_{iv}^{(b)}$ and variances $\hat{s}_{iv}^{2(b)}$; (iii) fitting ComBat model with the original data and obtaining scanner-specific means $\hat{\theta}_{iv}$ and variances $\hat{s}_{iv}^{2}$ and treating them as population parameters. We repeat this procedure $1,000$ times, and compute bias$^2$ and variance for scanner-specific means and variances for each candidate $r$. Finally, we compare mean squared error (bias$^2$+variance) to select the $r$ that achieves small bias$^2$ and variance simultaneously.

The bootstrap results for SPINS cortical thickness data are reported in the Supplementary Table 1 and 2. Based on this analysis, a value between $r = 5$mm and $r = 10$mm is a good choice, since both bias$^2$ and variance initially decrease and then increase, with turning points falling within the range of $r = 5$mm to $r = 10$mm. Therefore, selecting $r = 5$mm is reasonable according to the bootstrapping procedure.

| $r$ | 0mm | 5mm | 10mm | 20mm |
|---:|---:|---:|---:|---:|
| bias$^2$ | 106.43 | 67.56 | 72.66 | 73.50 |
| variance | 92.06 | 52.57 | 56.35 | 59.15 |

Supplementary Table 1: The bootstrapping bias$^2$ and variance of scanner-specific means on candidate $r$.

| $r$ | 0mm | 5mm | 10mm | 20mm |
|---:|---:|---:|---:|---:|
| bias$^2$ | 177.85 | 58.00 | 52.92 | 83.47 |
| variance | 172.35 | 54.02 | 45.38 | 78.49 |

Supplementary Table 2: The bootstrapping bias$^2$ and variance of scanner-specific variances on candidate $r$.

### S2 Spatial smoothing amplifies the covariance heterogeneity between scanners

For simplicity, suppose that a pair of imaging features is measured from two different scanners ($i = 1, 2,\ n_i = n$). We assume that $(y_{ij1}, y_{ij2})^\top$ follows spatial Gaussian process with mean zeros and scanner-specific variance-covariance structure,

$$\begin{pmatrix} y_{ij1} \\ y_{ij2} \end{pmatrix} \sim \mathcal{MVN}\left(\begin{pmatrix} 0 \\ 0 \end{pmatrix}, \begin{bmatrix} \tau_i^2 + \sigma_i^2 & \sigma_i^2\rho \\ \sigma_i^2\rho & \tau_i^2 + \sigma_i^2 \end{bmatrix}\right),$$

where $0 < \rho < 1$ is analogous to $\exp(-\phi \cdot d)$ or $\exp(-\phi \cdot d^2)$ in SAN. If we apply smoothing, consider weights $w_1 > 0$ and $w_2 > 0$ that are $w_1 + w_2 = 1$. We have $y^s_{ij1} = w_1 y_{ij1} + w_2 y_{ij2}$ and $y^s_{ij2} = w_2 y_{ij1} + w_1 y_{ij2}$ where

$$\begin{pmatrix} y^s_{ij1} \\ y^s_{ij2} \end{pmatrix} \sim \mathcal{MVN}\left(\begin{pmatrix} 0 \\ 0 \end{pmatrix}, \begin{bmatrix} (w_1^2 + w_2^2)(\tau_i^2 + \sigma_i^2) + 2w_1w_2\sigma_i^2\rho & (w_1^2 + w_2^2)\sigma_i^2\rho + 2w_1w_2(\tau_i^2 + \sigma_i^2) \\ (w_1^2 + w_2^2)\sigma_i^2\rho + 2w_1w_2(\tau_i^2 + \sigma_i^2) & (w_1^2 + w_2^2)(\tau_i^2 + \sigma_i^2) + 2w_1w_2\sigma_i^2\rho \end{bmatrix}\right).$$

To examine the covariance differences between scanners, we use the $F$ statistic formula by plugging the true parameters:

$$F = \frac{[\mathrm{E}(y_{1j1}y_{1j2}) - \mathrm{E}(y_{2j1}y_{2j2})]^2}{[\mathrm{Var}(y_{1j1}y_{1j2}) + \mathrm{Var}(y_{2j1}y_{2j2})]/n} = \frac{(\sigma_1^2\rho - \sigma_2^2\rho)^2}{[(\tau_1^2 + \sigma_1^2)^2 + \sigma_1^4\rho^2 + (\tau_2^2 + \sigma_2^2)^2 + \sigma_2^4\rho^2]/n},$$

which follows from $\mathrm{E}(y_{ij1}y_{ij2}) = \sigma_i^2\rho$ and $\mathrm{Var}(y_{ij2}y_{ij2}) = (\tau_i^2 + \sigma_i^2)^2 + \sigma_i^4\rho^2$. Similarly, the $F$ statistic for the smoothed data, denoted by $F^s$, is

$$\begin{aligned} F^s &= \frac{[\mathrm{E}(y^s_{1j1}y^s_{1j2}) - \mathrm{E}(y^s_{2j1}y^s_{2j2})]^2}{(\mathrm{Var}(y^s_{1j1}y^s_{1j2}) + \mathrm{Var}(y^s_{2j1}y^s_{2j2}))/n} \\ &= \frac{[\sigma_1^2\rho + w(\tau_1^2 + \sigma_1^2) - (\sigma_2^2\rho + w(\tau_2^2 + \sigma_2^2))]^2}{[((\tau_1^2 + \sigma_1^2) + w\sigma_1^2\rho)^2 + (\sigma_1^2\rho + w(\tau_1^2 + \sigma_1^2))^2 + ((\tau_2^2 + \sigma_2^2) + w\sigma_2^2\rho)^2 + (\sigma_2^2\rho + w(\tau_2^2 + \sigma_2^2))^2]/n}, \end{aligned}$$

where $w = \frac{2w_1w_2}{w_1^2 + w_2^2}$. If smoothed features show a larger covariance heterogeneity between scanners than unsmoothed features, the statistic $F^s$ of smoothed features will be larger than the unsmoothed $F$. When $\tau_i$ and $\sigma_i^2$ are monotonic (e.g., $\tau_1^2 > \tau_2^2$, then $\sigma_1^2 > \sigma_2^2$) we have $F^s > F$ because $F^s/F = A \times B$ where

$$\begin{aligned} A &= \left(w_1^2 + w_2^2 + \frac{2w_1w_2}{\rho}\left(1 + \frac{\tau_1^2 - \tau_2^2}{\sigma_1^2 - \sigma_2^2}\right)\right)^2 \\ B &= \frac{(\tau_1^2 + \sigma_1^2)^2 + \sigma_1^4\rho^2 + (\tau_2^2 + \sigma_2^2)^2 + \sigma_2^4\rho^2}{(w_1^2 + w_2^2)[((\tau_1^2 + \sigma_1^2) + w\sigma_1^2\rho)^2 + (\sigma_1^2\rho + w(\tau_1^2 + \sigma_1^2))^2 + ((\tau_2^2 + \sigma_2^2) + w\sigma_2^2\rho)^2 + (\sigma_2^2\rho + w(\tau_2^2 + \sigma_2^2))^2]}. \end{aligned}$$

Here, $A > 1$ follows from $2w_1w_2 < \frac{2w_1w_2}{\rho}\left(1 + \frac{\tau_1^2 - \tau_2^2}{\sigma_1^2 - \sigma_2^2}\right)$, and $B > 1$ is derived by using

$$((w_1^2 + w_2^2)(\tau_i^2 + \sigma_i^2) + 2w_1w_2\sigma_i^2\rho)^2 + ((w_1^2 + w_2^2)\sigma_i^2\rho + 2w_1w_2(\tau_i^2 + \sigma_i^2))^2 < (\tau_i^2 + \sigma_i^2)^2 + \sigma_i^4\rho^2.$$

### S3 The results of SAN+CovBat and SAN+RELIEF

We evaluate the performance of SAN+CovBat and SAN+RELIEF by conducting all the analyses parallelly. Using SAN as a benchmark, we summarize their results in Supplementary Figures 1-5 and Table 3. Overall, SAN+CovBat and SAN+RELIEF demonstrate performance comparable to SAN in mitigating inter-scanner biases in covariance. However, SAN slightly outperforms SAN+RELIEF and SAN+CovBat in terms of CASH scores. In Supplementary Figures 1 and 5, SAN achieves lower CASH scores at shorter distances. In Supplementary Figure 2, we also evaluate the effect size of the differences in CASH scores among these methods using Cohen's $d$ (with signs preserved), which is calculated as follows:

$$\text{Cohen's } d = \frac{\texttt{mean}(\text{CASH}_2) - \texttt{mean}(\text{CASH}_1)}{\texttt{sd}(\text{CASH}_2 - \text{CASH}_1)}$$

The calculated Cohen's $d$ values are 0.32 (SAN vs SAN+CovBat), 0.42 (SAN vs SAN+RELIEF), and 0.17 (SAN+CovBat vs SAN+RELIEF). These positive yet moderate Cohen's $d$ values indicate that SAN has slightly lower CASH scores compared to SAN+RELIEF and SAN+CovBat. This suggests that spatial covariance explains most of the inter-scanner covariance heterogeneity of cortical thickness in SPINS study, indicating that SAN alone is effective in addressing this issue.

The supplementary figures 1-5 and supplementary table 3 reproduces the results in the main manuscript, including SAN, SAN+RELIEF, and SAN+CovBat only for comparison purposes.

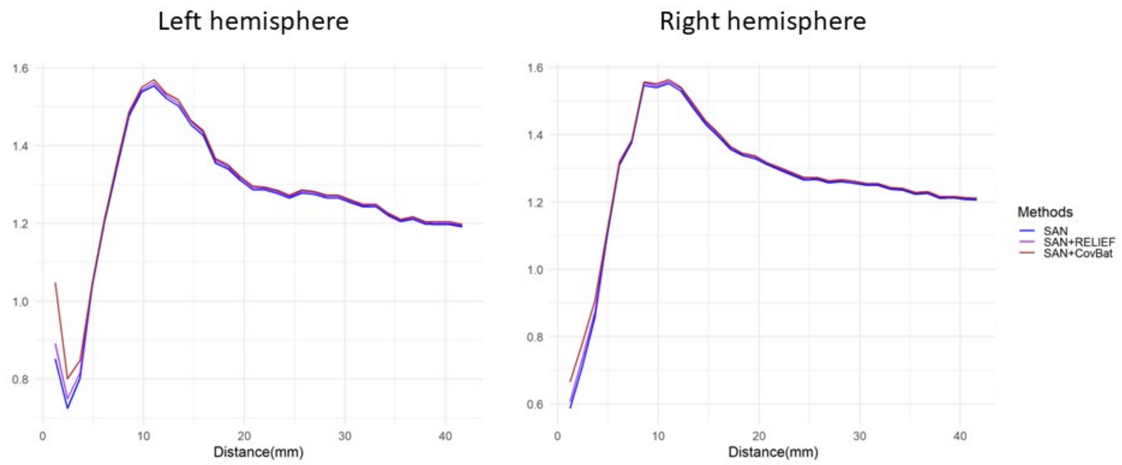


Supplementary Figure 1: Summary of CASH scores obtained from SAN related methods.

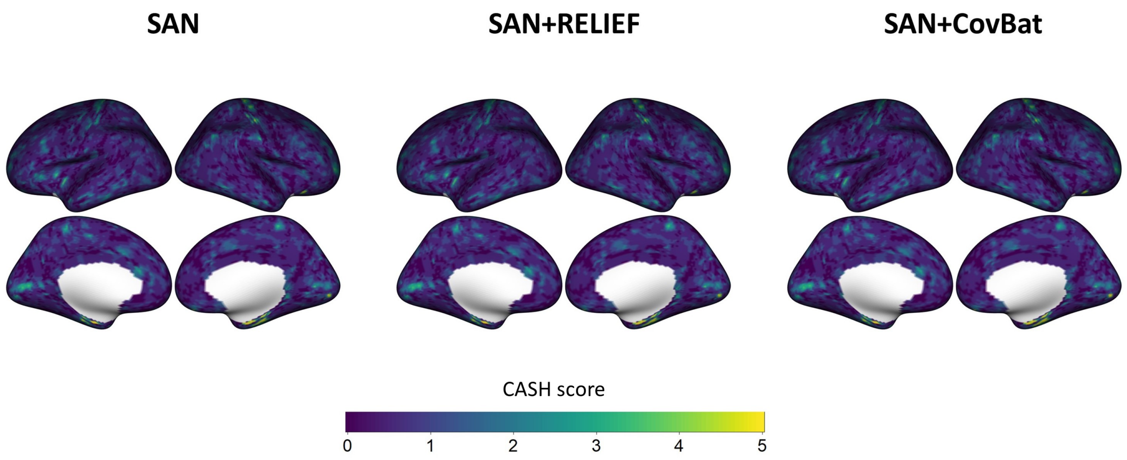


Supplementary Figure 2: CASH score brain maps obtained from SAN-related methods.

| Measure | CASH | Accuracy | AUC |
|---|---|---|---|
| SAN | 2.351 | 0.608 | 0.601 |
| SAN+RELIEF | 2.352 | 0.608 | 0.601 |
| SAN+CovBat | 2.360 | 0.611 | 0.603 |

Supplementary Table 3: The summary of CASH score, the accuracy and AUC of predicting scanners. The smallest values for SAN related methods are bolded respectively.

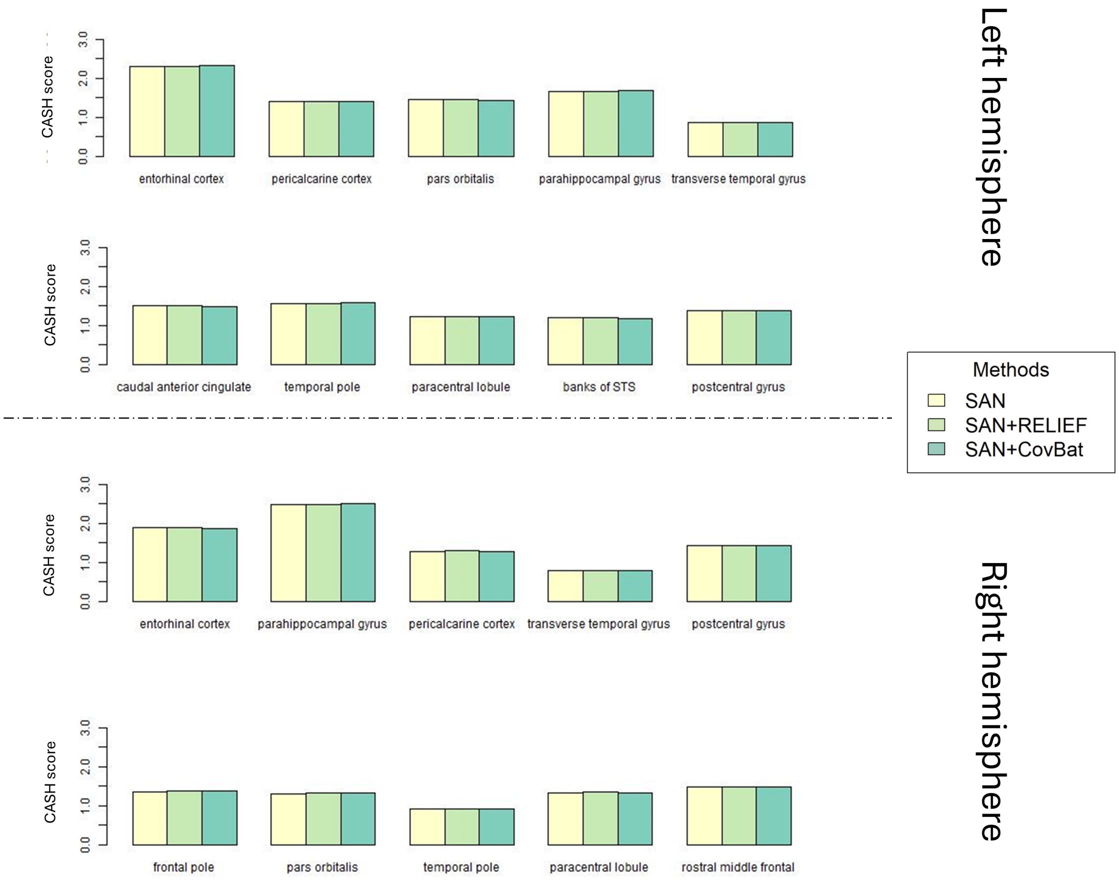


Supplementary Figure 3: CASH scores obtained from SAN related harmonization methods for the top 10 regions in each hemisphere, selected based on their highest CASH scores in the raw data.

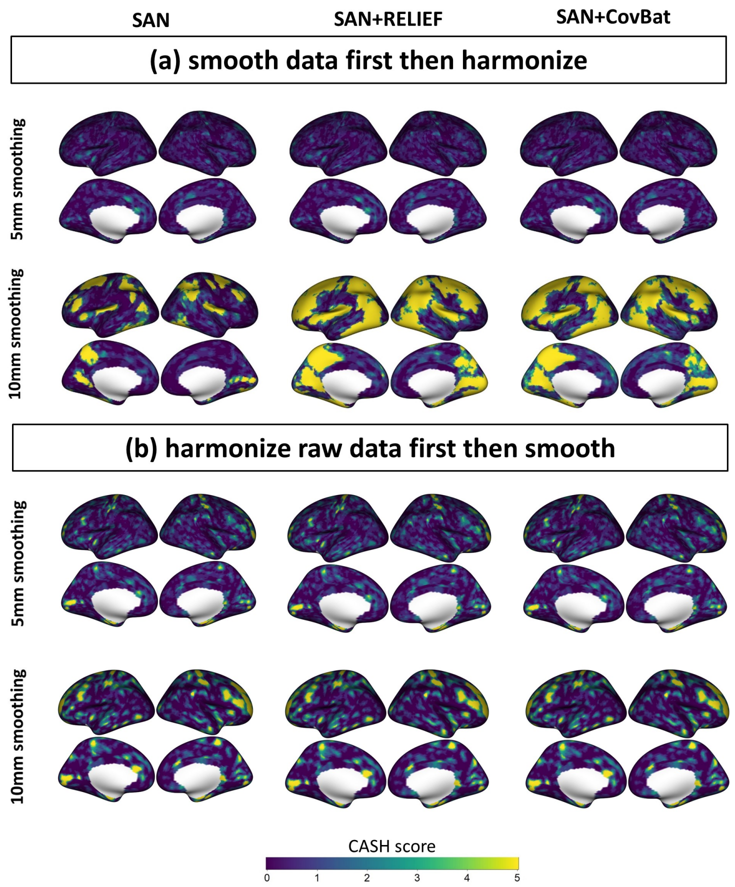


Supplementary Figure 4: CASH score brain maps on smoothed data obtained from SAN related methods.

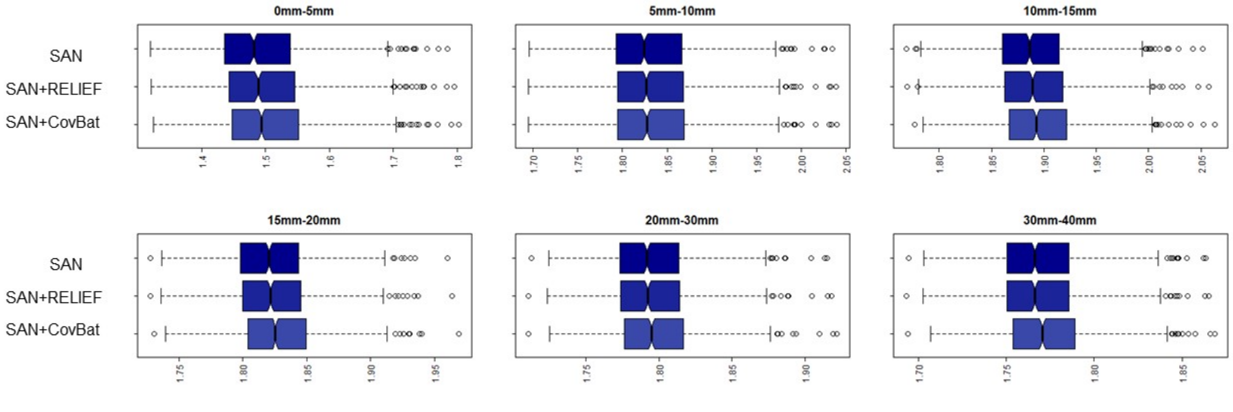


Supplementary Figure 5: Box plots illustrating CASH scores for SAN related methods across different intervals in the Bootstrapping results.

### S4 QDA for scanner prediction at the vertex level

QDA requires n>V (the sample size is larger than the number of features), which is not applicable to all vertex-level features due to the high dimensionality. To circumvent this limitation, we selected 10 representative clusters for each hemisphere that exhibit high CASH score in the raw data. This selection allows us to apply QDA on these clusters.

The selection procedure is as follows:

1. Arrange the vertex CASH score in the raw data in descending order.
2. Select the vertex with the highest CASH score and its adjacent vertices within a 10mm radius to form the first cluster.
3. For each subsequent cluster, choose the vertex with the highest CASH score among those whose distance from previously selected vertices exceeds 10mm to prevent overlapping clusters.
4. Repeat Step 3 iteratively to form all clusters.

These clusters are not only selected for their pronounced covariance scanner effects but also for their representativeness across different regions. This allows for a clearer comparison of performance differences among harmonization methods. For each vertex cluster set, we use leave-one-out cross-validation to calculate the average accuracy as well as ROC curve's area under the curve (AUC) for each harmonized dataset after regressing out covariate effects. The scatterplots of accuracy and AUC between 7 harmonization methods are shown in Supplementary Figure 6 and 7.

From Supplementary Figure 6 and 7, we observe that SAN+CovBat performs best in achieving lower accuracy and AUC for predicting scanners, as evidenced by all its fitted lines deviating from the diagonal lines and toward the direction where other methods' accuracy and AUC are larger. This underscores its effectiveness in comprehensively addressing inter-scanner covariance heterogeneity. SAN, CovBat, SAN+RELIEF, and ComBat perform worse than SAN+CovBat, but they all show much improvement in impairing detection of scanner compared to raw data. RELIEF also shows some improvement from raw data, but the differences are not as large as the other harmonization methods.

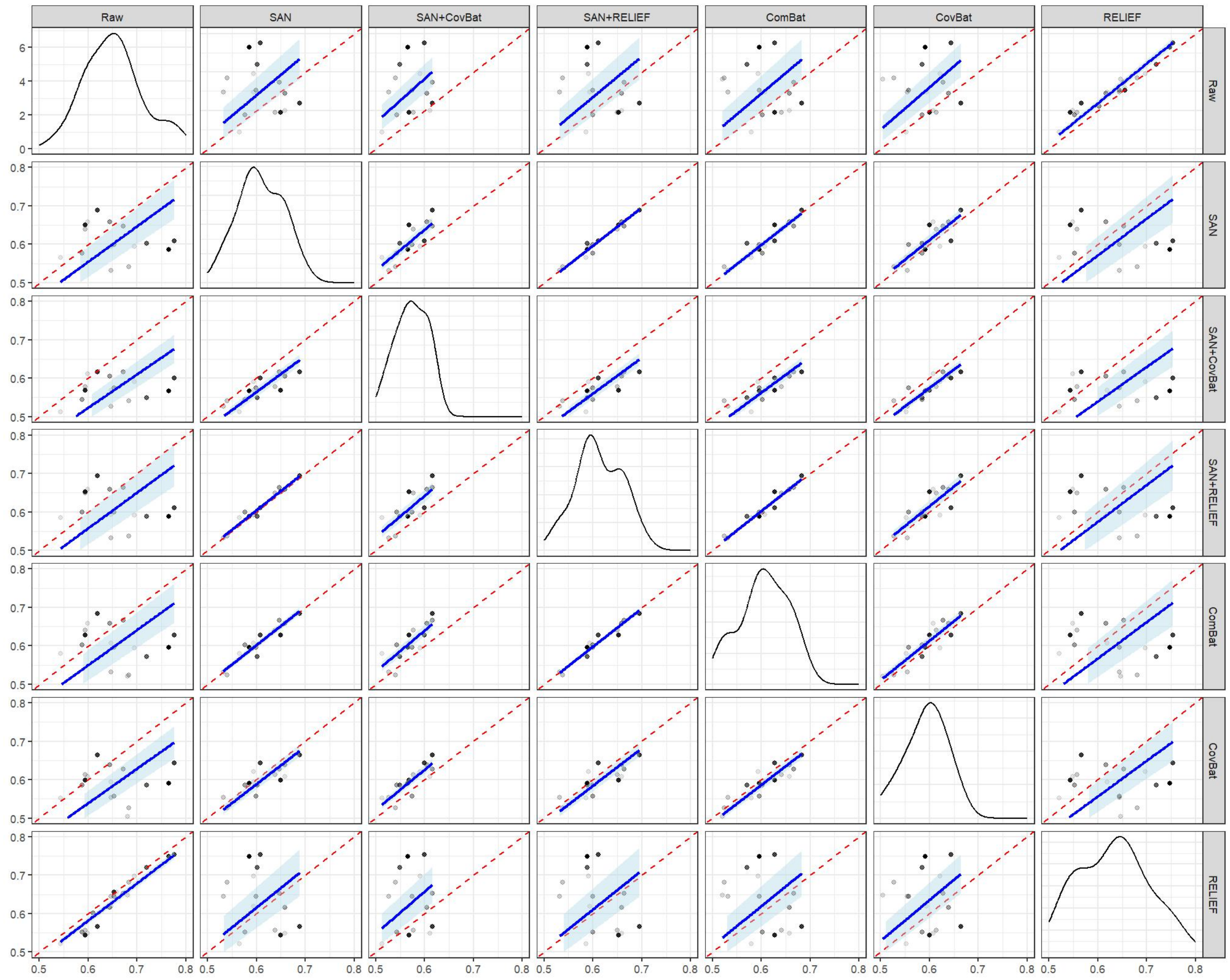


Supplementary Figure 6: We used QDA as a classifier and obtained the predictions on 20 cluster sets. The non-diagonal panels display scatterplots of accuracy for scanner predictions. Density estimations of accuracy are presented in the diagonal panels. In the scatterplots, transparency levels correspond to the CASH score, with darker shades indicating higher values. The red dashed lines represent diagonal lines (45 degree). The blue lines are fitted lines with 95% confidence intervals from a regression model without intercepts .

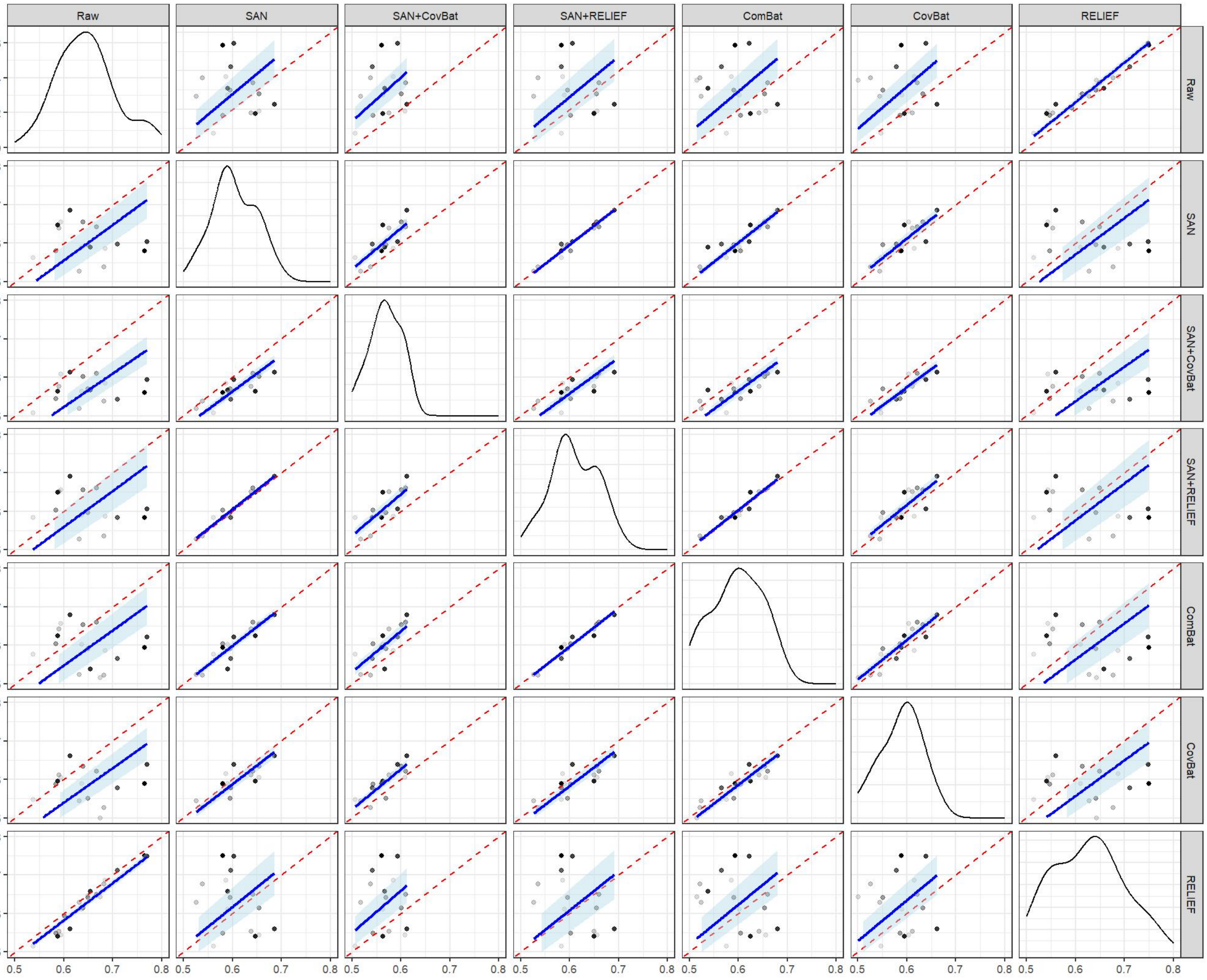


Supplementary Figure 7: We used QDA as a classifier and obtained the predictions on 20 cluster sets. The non-diagonal panels display scatterplots of Area Under the Curve (AUC) for scanner predictions. Density estimations of Area Under the Curve (AUC) are presented in the diagonal panels. In the scatterplots, transparency levels correspond to the CASH score, with darker shades indicating higher values. The red dashed lines represent diagonal lines (45 degree). The blue lines are fitted lines with 95% confidence intervals from a regression model without intercepts .

### S5 Within-ROI results for the right hemisphere

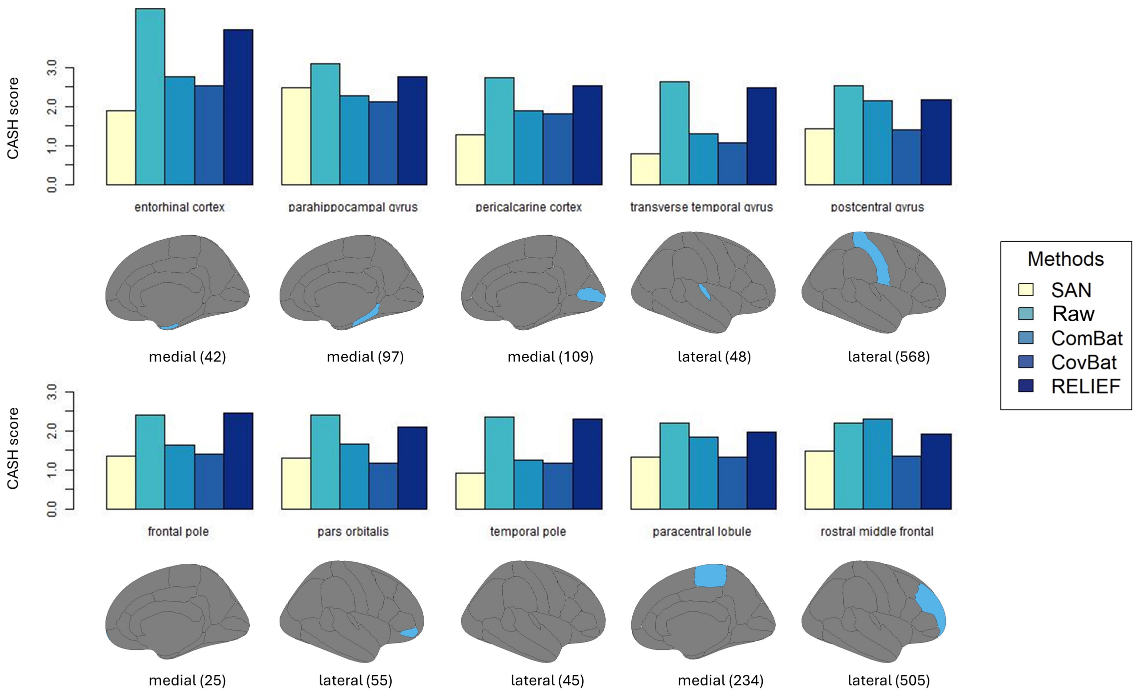


Supplementary Figure 8: ROI-level CASH scores from different harmonization methods. The top 10 regions in the right hemisphere are selected based on their highest CASH scores from the raw data. Below the corresponding bar plots, diagrams of the ROI complete with vertex counts are displayed.
